# Morphological plasticity in *Chlamydomonas reinhardtii* and acclimation to micropollutant stress

**DOI:** 10.1101/2020.08.29.273557

**Authors:** Giulia Cheloni, Vera I. Slaveykova

**Author notes:** The authors declare no competing financial interest.

## Abstract

Phytoplankton are characterized by a great phenotypic plasticity and amazing morphological variability, both playing a primary role in the acclimation to changing environments. However, there is a knowledge gap concerning the role of algal morphological plasticity in stress responses and acclimation to micropollutants. The present study aims examining the palmelloid colony formation of the green alga Chlamydomonas reinhardtii upon micropollutants exposure.

Cells were exposed to four micropollutants (MPs) with different modes of action (copper, cadmium, PFOS and paraquat) for a duration of 72h. Effects of MPs on palmelloid formation, growth and physiological traits (chlorophyll fluorescence, membrane integrity and oxidative stress) were monitored via flow cytometry and fluorescence microscopy. Palmelloid formation was observed upon treatment with the four micropollutants. Number of palmelloid colonies and their size were dependent on MP concentration and exposure duration. Cells reverted to their unicellular lifestyle when colonies were harvested and inoculated in fresh medium indicating that palmelloid formation is a plastic response to micropollutants. No physiological effects of these compounds were observed in cells forming palmelloids and palmelloid colonies accumulated lower Cd concentration than unicellular *C. reinhardtii* suggesting that colony formation protects the cells form MPs exposure. The results show that colony formation in *Chlamydomonas reinhardtii* is a stress response strategy activated to face sub-lethal micropollutant concentrations.

**HIGHLIGHTS:**

- Sub-lethal concentrations of micropollutants (MPs) induce palmelloid formation in *C. reinhardtii*
- Morphological changes are not associated to adverse effects on algal cells
- Palmelloid formation is transitory, cells revert to unicellular lifestyle in the absence of MPs
- Cells within large colonies experience lower Cd exposure than unicellular *C. reinhardtii*
- Palmelloid formation is a morphological stress response that plays a role in cells acclimation to MPs

## INTRODUCTION

One of the current challenges in environmental toxicology and risk assessment of micropollutants is related with the broad range of sensitivity that different species of the same group might have toward one or multiple micropollutants (MPs) (Segner et al., 2014). Such variation in sensitivity is mainly associated to the ability and strategies that different organisms might have to respond and acclimate to the stressors. In the last years, phytoplankton environmental toxicology investigations focused mainly on adverse effects induced by MPs, while mechanisms activated to respond and maintain cellular homeostasis at sub-toxic level exposure are still poorly understood. Phenotypic plasticity responses are associated with acclimation of the organism to MPs and are reversible when the stress is released. Such stress responses in phytoplankton were mostly investigated at molecular and physiological level (Mendez-Alvarez et al., 1999) while changes in morphological traits and behavior are rarely taken into account.

Some green algal taxa might undergo morphological modifications in presence of environmental constraints. Colony formation in green algae were reported to occur in response to predators (Lürling, 2009; Van Donk et al., 2010) and to low nutrients availability (Shubert et al., 2014; Trainor, 1970) as well as to changes in water salinity (Keerthi et al., 2016). However, a limited knowledge is available on algal morphological plasticity in response to micropollutant stress, despite the recognized importance of morphological traits in shaping phytoplankton natural communities exposed to toxic chemical pollutants (Fischer et al., 2013).

In this study, we investigated colony formation of the green alga *Chlamydomonas reinhardtii* upon exposure to micropollutants. *Chlamydomonas* species are known to form micro-colonies of cells dubbed palmelloids, however very little is known about its colonial lifestyle and palmelloid condition. Formation of palmelloids or groups of nonflagelled cells is not uncommon in laboratory culture of this alga (Harris, 2009); and the mechanisms that drive the transition from unicellular to colonial lifestyle are poorly understood. Palmelloid formation is generally associated with the adverse environmental conditions, such as presence of predators (Herron et al., 2019; Lurling and Beekman, 2006), salt stress (Khona et al., 2016; Takouridis et al., 2015) and organic acids induced stress (Iwasa and Murakami, 1969). Possible palmelloid formation in presence of micropollutants was rarely reported (Goff et al., 2013; Jamers and De Coen, 2010) despite the extended use of *Chlamydomonas* strain in ecotoxicological studies. However, long-term changes in algal cell morphology are never taken into consideration as toxicity endpoints. This study thus aims to provide new insight into the morphological plasticity of *C. reinhardtii* in response to MPs exposure and to highlight the role of such cellular response in algal acclimation to MPs-induced stress.

## MATERIAL AND METHODS

### Strain and culturing system

*Chlamydomonas reinhardtii* CPCC11 (Canadian Phycological Culture Centrer, Department of Biology, University of Waterloo, Canada) was axenically grown in a modified HSM medium (Lavoie et al., 2009). Cultures were incubated in a specialized incubator (Fitotron, Weiss-technique, Switzerlan) at 20°C, under 85 μmols photons m^-2^ s^-1^ with a 16:8 light cycle under rotary shaking at 110 rpm. Glassware were soaked for at least 24 h in 5% v/v with HNO_3_ and rinsed at least three times with MilliQ water. Reagents were analytical grade. Stock solutions and media were prepared with ultrapure MilliQ water (>18.2Ω MilliQDirect system, Merck Millipore, Darmstadt, Germany).

### Micropollutant treatments and biological endpoints

Four micropollutants, two organic compounds, Paraquat and perfluorooctanesulfonic acid (PFOS), and two metals, Cd and Cu, were selected since characterized by different modes of toxic action. Paraquat is an herbicide that acts on photosystem I and promotes the formation of reactive oxygen species by interfering with electron transfer (Jamers and De Coen, 2010). PFOS is a fluorosurfactants widely used in industrial processes, its mode of toxic action in phytoplankton is still unknown. In eukaryotic cells is known to affect fatty acid metabolism and membrane stability (U.S. Environmental Protection Agency). Copper and cadmium were selected as representative of redox active essential metal and redox inactive non-essential metal, respectively, which are able to affect *Chlamydomonas* cells (Szivak et al., 2009).

### Exposure condition

Cultures were maintained in exponential growth for at least 2 weeks before starting tests with MPs. *C. reinhardtii* cultures with density of 2.5×10^5^ cell mL^-1^ were exposed to increasing MP concentrations in 100 mL Erlenmeyer flasks and incubated for 72h in specialized incubators under the conditions previously specified. PFOS and paraquat (Sigma–Aldrich, Buchs, Switzerland) stock solutions were prepared in MilliQ water, while Cd(NO_3_)_2_ and CuSO_4_ standard solutions were purchased (AAS, Sigma–Aldrich, Buchs, Switzerland) and diluted before utilization. Exposure concentration ranges for PFOS, paraquat, Cu and Cd were 50-750 μM, 25nM-2.5 μM, 0.1-2.5 μM and 0.01-2.4 μM, respectively. For Cd and Cu the exposure concentrations are expressed as free metal ions (total [Cu] 0.8-5 μM and total [Cd] 0.4-3.5 μM). The initial free Cu and Cd ion concentrations ([Cu^2+^], [Cd^2+^]) in the exposure medium were calculated by visual MINTEQ version 3.0 (Gustafsson, 2010) based on the total ion concentration measured by inductively coupled plasma mass spectrometry (ICP-MS, 7700x Agilent Technologies).

### Effect of MPs on algal morphology

Possible effect of MPs on morphological and physiological traits of *C. reinhardtii* were investigated. Algal growth and morphological changes were followed via direct microscopic observation (BX61, Olympus, Volketswil, Switzerland) of *C. reinhardtii* cultures. Samples were collected at 2, 24, 48 and 72 h, fixed with 0.25% glutaraldehyde and stored in the dark at 4°C until imaging. For each sample replicate at least 200 particles were imaged.

Three morphotypes were identified (**Figure 1**): (i) unicellular healthy, mostly flagelled cells with ellipsoid shape and a large green chloroplast that occupies the whole cell; (ii) palmelloid microcolonies of 2 to 16 cells; (iii) unicellular damaged cells with a round shape and smaller chloroplast of pale green color. Changes in the size and biovolume were investigated using a Coulter Counter Multisizer II (Beckman Coulter, Nyon, Switzerland) with a 50 μm orifice tube.

**Figure 1:**
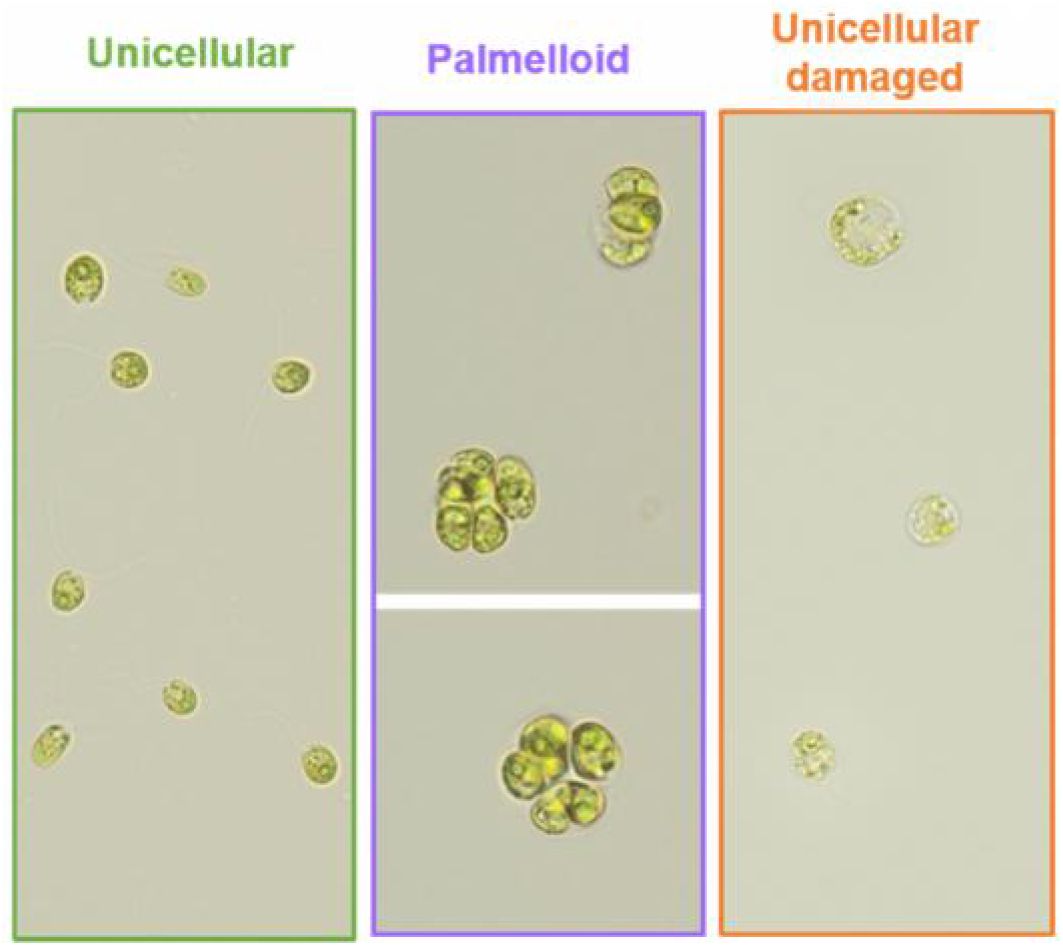
Changes in *C. reinhardtii* morphology were monitored by microscopy and three morphotypes were identified in the presence of MPs: (i) unicellular healthy cells; (ii) Palmelloid micro-colonies; (iii) unicellular damaged cells.

### Reversibility of palmelloid formation

To test further whether palmelloid formation is a reversible process, *C. reinhardtii* exposed to 1.5 μM [Cd^2+^] and 500 μM [PFOS] for 72h, that presented predominant palmelloid morphology, were harvested by centrifugation. Half of the harvested palmelloid colonies were re-suspended in exposure medium containing the same MP concentration while the other half was re-suspended in exposure medium with no Cd or PFOS. Morphological changes were followed via direct microscopic observation (BX61, Olympus, Volketswil, Switzerland).

### Phisioloigcal effect of the MPs on algae with different morphology

Based on the morphological observation, three different sets of experiments were performed corresponding to: (i) no effect on algal morphology, (ii) palmelloid formation and (iii) cellular damages. In each experimental set, MPs-induced changes in physiological traits such as chlorophyll fluorescence, membrane permeability and oxidative stress were followed by flow cytometry. Measurements were performed by BD Accuri C6 (Accuri cytometers Inc., Michigan) flow cytometer equipped with a blue light (488 nm) excitation laser and three detectors for determination of green (530±15 nm), yellow (585± 20 nm) and red (670± 25 nm) fluorescence. Algal suspensions were collected after 48h-exposure and passed through FCM using medium flow rate. The threshold was set to 20000 events. Data acquisition and analysis was performed with BD Accuri C6 Software 264.15. A gating strategy was developed to discriminate between *C. reinhardtii* unicellular and palmelloid particles (**Figure S1**). Two distinct particle populations were defined for unicellular algae and palmelloid colonies based on the forward scattering (associated to size) and side scattering (associated to granularity) properties of the particles. Bleaching of chlorophyll fluorescence were determined by following the variations in the chlorophyll autofluorescence of the algal populations. The percentage of cells with oxidative stress and with damaged membranes were determined using the fluorescent probes CellROX®Green (Life Technologies Europe B.V., Zug, Switzerland) and propidium iodide (PI, Sigma–Aldrich, Buchs, Switzerland), following the previously developed methodology (Cheloni et al., 2014) and applying the gating strategy shown in Figure S1.

*Metal accumulation*. Cd accumulation by *C. reinhardtii* was determined daily during the process of palmelloid formation. To this end *C. reinhardtii* was isolated form the exposure medium by gentle centrifugation at 3500 × g for 5 min. The supernatant was collected and acidified with HNO_3_ (Suprapur, Merck, Darmstadt, Germany). Metal content was measured, after appropriate sample dilution, by ICP-MS. The pellet was re-suspended in the medium supplemented with 1 mM EDTA (Sigma–Aldrich, Buchs, Switzerland) for 10 min and centrifuged. The EDTA-washed pellets were digested with 65% HNO_3_ (Suprapur, Merck, Darmstadt, Germany) at 90°C for 2 h to determine the EDTA non-extractable metal content. Samples were measured, after appropriate dilution by ICP-MS. In order to verify the influence of palmelloid colony size on Cd uptake, palmelloids obtained upon 24, 48 and 72h exposure to PFOS were collected by gentle centrifugation at 3500 × g for 5 min, rinsed with M-HSM medium and resuspended in M-HSM medium containing 1.5 μM [Cd^2+^]. Samples were collected after 2 h exposure and treated as described above in order to measure the EDTA extractable metal adsorbed on *C. reinhardtii* surface and the EDTA non-extractable metal content.

*Statistical analysis* was performed comparing the results obtained for samples treated with MPs to those obtained for untreated control samples using a Mann-Whitney Rank Sum Test. All statistical analysis was performed using the software Sigma Plot version 12.5 (Systat Software Inc., San Jose, CA).

## RESULTS AND DISCUSSIONS

### Morphological changes upon MPs exposure

Palmelloid formation was observed upon 48h-exposure to all the MPs tested and depended on the MP concentration and exposure time (**Figure 2**). Interestingly, the range of concentrations able to induce palmelloid formation changed considerably among micropollutants. For instance, palmelloid formation was observed only during exposure to 1.3 and 1.6 μM [Cu^2+^], whereas exposure to Cd concentration higher than 0.4 μM Cd^2+^ induced palmelloid formation in the majority of the exposed cells. Similar results were obtained upon exposure to PFOS, but only at the highest concentrations tested. Opposite to the observation for Cd, Cu and PFOS, exposure to paraquat led to the formation of palmelloids in less than the half of the algal population. At higher MPs concentrations such as 1.9 μM [Cu^2+^], 2.4 μM [Cd^2+^] and 0.75 μM [paraquat], a decrease in the number of cells in palmelloids was observed. At these concentrations, cells with evident damage were also present in the cultures. The observed decrease of palmelloid colonies and the increase of damaged cells indicate that MPs exposure either induces palmelloid formation or damaged morphotypes. No palmelloid colonies with evident cellular damages were detected during microscopic investigation. It is important to note that palmelloid colony formation was observed during exposure to all the tested MPs, indicating that this morphological plasticity is a common cellular response rather than a specific response related to the mode of action of a given stressor. This observation is in line with previous reports on palmelloid formation of *C. reinhardtii* exposed to different biotic and abiotic stressors (Goff et al., 2013; Herron et al., 2019; Iwasa and Murakami, 1969; Jamers and De Coen, 2010; Khona et al., 2016; Lurling and Beekman, 2006; Takouridis et al., 2015).

**Figure 2:**
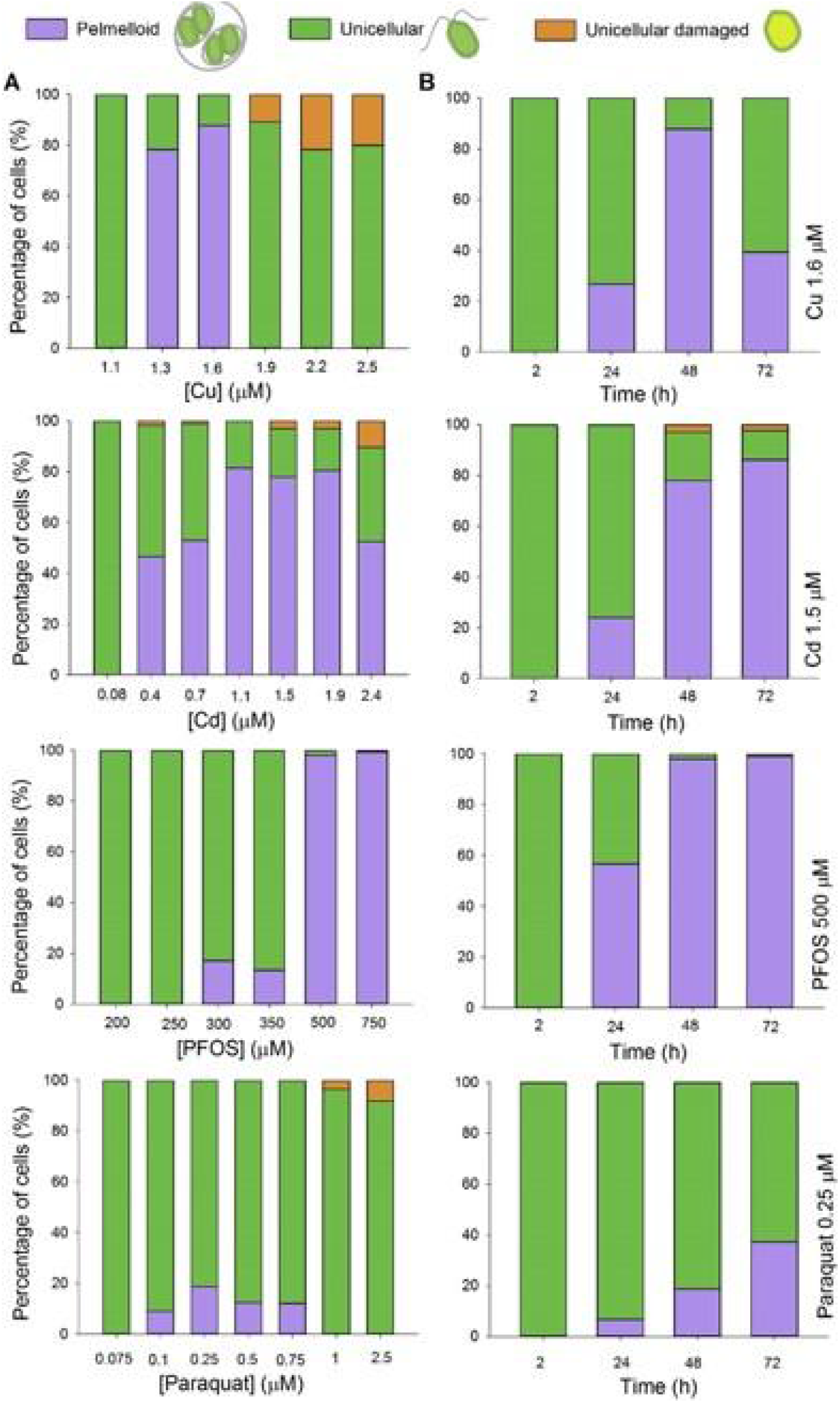
Changes in cell morphology as a function of micropollutants concentration after 48h exposure **(A)** or exposure time for a given micropollutant concentration **(B)**. Stacked bars indicate percentage of cells forming palmelloid colonies (violet bars) or that had unicellular (green bars) or damaged unicellular (orange bars) morphology. Data for all the tested concentrations and associated standard deviations are available in Table S1.

Differences in the time course of palmelloid formation were observed among MPs treatments as a function of exposure time. In 0.25 μM paraquat treatment the highest yield of palmelloid formation was reached after 72 h, while the peak was observed after 48h exposure to 1.5 μM [Cd^2+^], 1.6 μM [Cu^2+^] and 500 μM [PFOS]. Cd and PFOS had similar patterns of palmelloid formation with almost all of the cells in palmelloids for a given concentration after 48h, and palmelloid condition was found to be stable through time. In contrast, palmelloid formation during Cu exposure was transient (Figure 2). In the 1.6 μM [Cu^2+^] treatment, the maximum percentage of palmelloids was observed after 48h exposure, followed by a decrease in the percentage of cells forming palmelloids after 72h. Microscopic observations and particle count data revealed that the increase of the percentage flagelled cells at 72h exposure was due to the release of cells from the palmelloid colonies rather than to the growth of the unicellular cells during 48h exposure. Such reversibility of the colonies to unicellular lifestyle might be associated with a decrease of the [Cu^2+^] and thus bioavailability after 48h incubation. Indeed, a decrease of dissolved Cu over exposure time was found (Figure S2). What is more the bioavailable fraction of Cu after 48h incubation might be considerably lowered due to the release of biological ligands in the exposure medium (Kola et al., 2004).

### Reversibility of palmelloid formation

Palmelloid colonies resuspended in exposure medium in presence of Cd and PFOS kept palmelloid morphology (**Figure 3**) and colony size became larger. When resuspended in exposure medium in the absence of MPs, half of the *C. reinhardtii* cells remained in palmelloid after 24h and only few colonies were present after 48h (**Figure 3**). These results indicate that in the absence of MPs, *C. reinhardtii* are capable to revert to their unicellular lifestyle and confirm that the process of palmelloid formation is associated to the presence of MPs in the ambient medium. Despite the different modes of action of Cd and PFOS, similar patterns of colony formation and disappearance were observed, confirming that palmelloid formation is a transitory process that is not specific to the MP. Reversibility of palmelloid formation was previously observed upon salt stress (Khona et al., 2016) and its transitory characteristic suggests that this cellular process might be a strategy to respond to stress pulses and micropollutants bursts.

**Figure 3:**
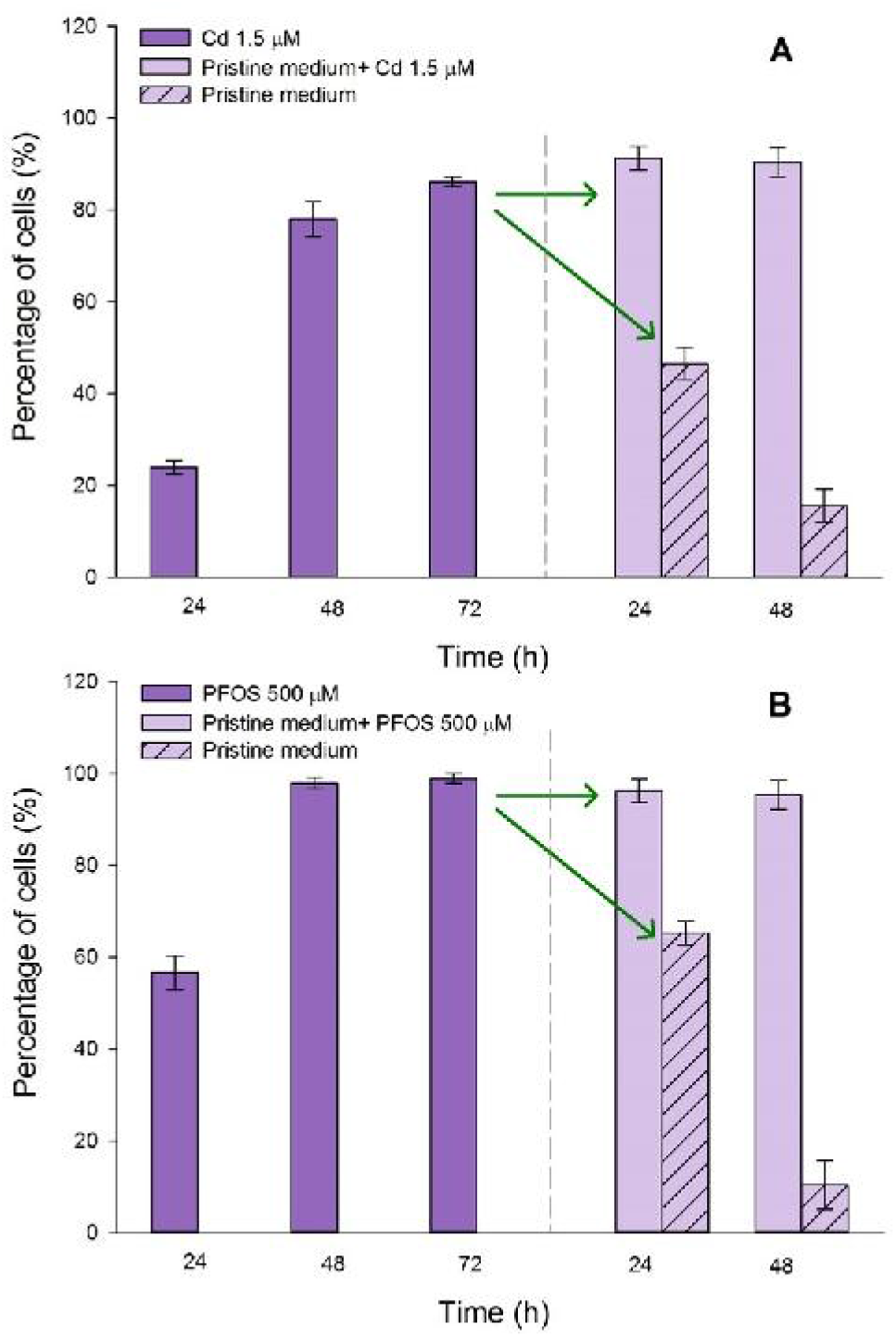
Percentage of cells that are in palmelloid colonies upon exposure to Cd **(A)** and PFOS **(B)**. After 72h incubation, palmelloid colonies were harvested and re-suspended in medium with no MPs (striped bars) and medium with Cd or PFOS (empty bars). Values are average ± standard deviations.

### C. reinhardtii growth and palmelloid formation

Microscopic observation of palmelloid colonies indicated that the number of cells present within each colony increases through time. For this reason *C. reinhardtii* growth was evaluated in two ways, by measuring the algal particle density (unicellular or palmelloid) via flow cytometry, and by counting the number of cells (unicellular or within the palmelloid colony) using microscopy. Data of algal density measured via FCM and direct cell count overlapped in unicellular *C. reinhardtii* cultures. By contrast, significant differences between particle number and cell number were observed in cultures where palmelloid colonies were abundant. Exposure to 1.6 μM [Cu^2+^] resulted in formation of considerable amounts of palmelloid colonies after 48 hours exposure (Figure 2) and this is reflected in the significant difference between the algal particle density and the cell density as well as in the increase in particle size (Figure 4). Such differences were no longer significant after 72h incubation when the cells reverted to the unicellular life style.

**Figure 4:**
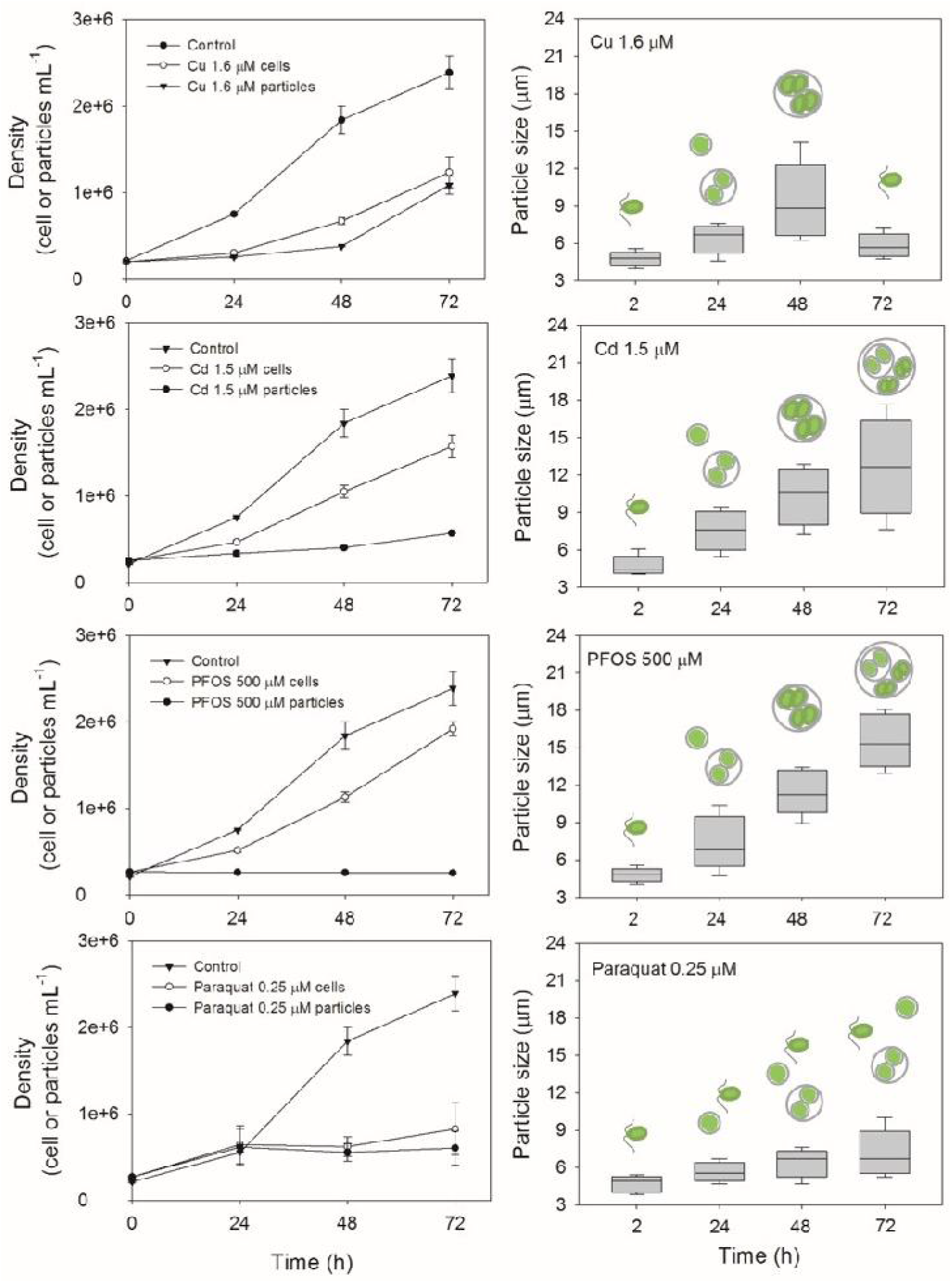
*C. reinhardtii* growth during palmelloid formation. On the left plots that show the changes in particle/cells density as a function of time for untreated cultures and cultures exposed to Cu, Cd, PFOS and paraquat concentrations able to induce palmelloid formation. Particles number was determined via flow cytometer counting while cell number was determined via direct microscope counting and takes into account the number of cells within palmelloid colonies. Values are the average of at least three replicates ± standard deviation. On the right side plots that show the changes in *C. reinhardtii* particle volume as a function of time upon exposure to Cd, Cu, PFOS and paraquat concentrations that induce palmelloid formation. Whiskers represent the 10^th^ and the 90^th^ percentiles.

No significant changes in the particle numbers over time was observed upon exposure to 1.5 μM [Cd^2+^] and 500 μM [PFOS] (Figure 4). Whereas, under these exposure conditions, the number of cells increased significantly indicating that cells keep growing and dividing within each palmelloid colony. The increase in the number of cells within each colony affects particle size and a significant increase of the median particle size values as a function of time was found under these conditions (Figure 4). Cellular mechanisms of palmelloid formation were not yet investigated but palmelloid formation seems to be associated with cellular growth. Electron microscopy observation of *Chlamydomonas* palmelloids revealed that cells inside the palmelloid are enveloped in multiple layers of cell wall (Iwasa and Murakami, 1969; Khona et al., 2016). For this reason, palmelloid formation is intuitively associated more to the retention of the daughter cells within the mother cell wall after cell division than to abnormal cell proliferation or aggregation of single cells (Harris, 2009).

No significant difference between particle and cell numbers was observed in the 0.25 μM paraquat treatment, which inhibited considerably *C. reinhardtii* growth and consequently palmelloid formation (Figure 4). The observed increase in particle size after 72h exposure to 0.25 μM paraquat was in part due to the formation of palmelloids and in part associated to the presence of large single cells of *Chlamydomonas reinhardtii*. The results of the present study are in accordance with what was previously reported for *C. reinhardtii* exposure to paraquat with growth inhibition at comparable concentrations (EC50 0.26 μM) and increase of size after long-term exposure (Jamers and De Coen, 2010).

Overall, the results obtained so far suggested that palmelloid formation occurred at sub-lethal MPs concentrations, and that cells kept growing and dividing within the palmelloid colony. However, the question whether the exposure to MPs induces only a shift from unicellular to colonial lifestyle or the physiology of palmelloid colonies is altered remains to be answered.

### Physiological characteristics of palmelloid colonies and unicellular damaged cells

To verify whether MPs induce physiological effects on palmelloid colonies we investigated possible changes in chlorophyll fluorescence, oxidative stress and membrane damage. Such toxicity endpoints are commonly investigated to evaluate MPs impact on *C. reinhardtii* (Cheloni et al., 2014; Nestler et al., 2012; Szivak et al., 2009). The effects on unicellular and palmelloid colonies in cultures were a mixture of the two population was present were followed by FCM (Figure 5). Algal particles were first separated in two populations (unicellular or palmelloid) using a gating strategy based on forward and side scatter values of the algal particles. For each population we could investigate in parallel possible changes in chlorophyll fluorescence, membrane damage and oxidative stress (Figure 5).

**Figure 5:**
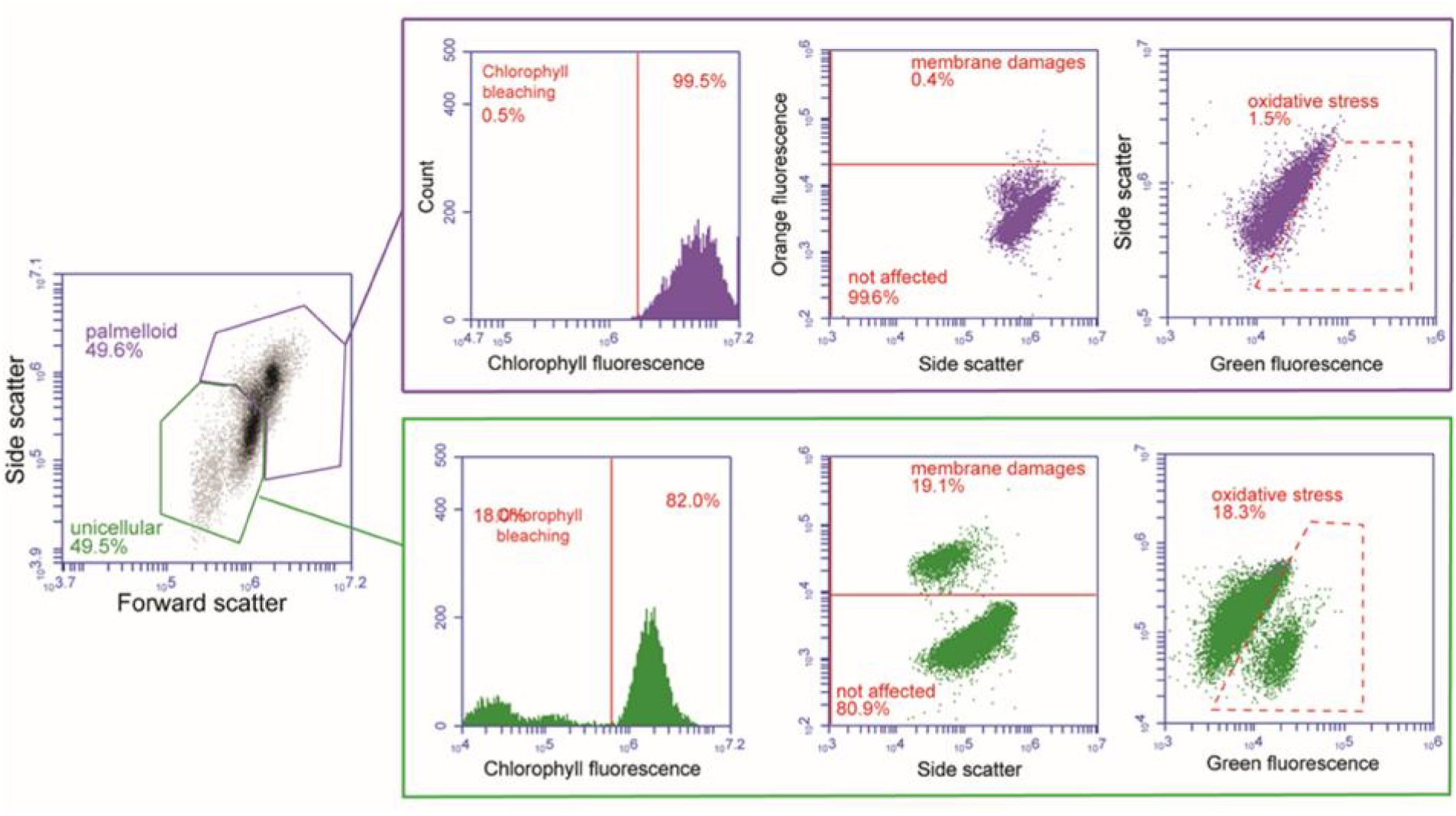
Example of the flow cytometry gating procedure that enables the discrimination between unicellular and palmelloid particle populations. Physiological effects (chlorophyll bleaching, membrane damage and oxidative stress) could be monitored in parallel for each of the two populations. Data obtained for a *C. reinhardtii* culture after 48h exposure to 1.6 μM [Cd^2+^].

No MPs-induced alteration on the chlorophyll fluorescence, oxidative stress and membrane damages were found under exposure conditions corresponding to the lack of morphological changes (unicellular culture) (Table 1). For the cultures characterized by mixed populations consisting of single cells and palmelloids (exposure to 0.25 μM [paraquat], 1.6 μM [Cu^2+^], 1.5 μM [Cd^2+^] or 500 μM [PFOS]), the unicellular population exhibited chlorophyll bleaching, membrane damage and oxidative stress, whereas palmelloids were unaffected.

Exposure to the highest concentration of Cu or paraquat resulted in strong effects for the tested endpoints in almost the totality of the unicellular algae. Interestingly, the FCM highlighted that cells within the same population might respond differently to the presence of the MPs. Such heterogeneity in the *C. reinhardtii* sensitivity towards MPs was previously reported (Cheloni et al., 2014; Szivak et al., 2009) highlighting that only part of the algal population is affected by the MPs despite the fact that all of the cells experienced the same exposure conditions.

The present results showed that MPs at specific concentration might induce morphological changes and palmelloid formation but no adverse effects on algal cells. The absence of stress in palmelloid colonies indicates that with this morphological response cells temporarily acclimate to presence of MPs. Such transition from unicellular to colonial lifestyle is transitory and cells revert to the unicellular flagelled form in the absence of stressors.

Palmelloid formation might have adaptive role when cultures are exposed chronically to predator pressure (Herron et al., 2019). Unicellular *C. reinhardtii* cells co-cultured with the predator *Paramecium tetraurelia* for 50 weeks evolved into multicellular forms. Such multicellularity conferred an effective defence against predation by rotifers (Herron et al., 2019) indicating that morphological changes might have important roles not only in acclimation but also in adaptive responses to environmental stressors. The investigation of *Chlamydomonas* morphological plasticity might be of particular relevance for the evaluation of the impact of chronic exposure of natural communities to low concentrations of MPs.

Overall, the results demonstrated that at the sub-toxic concentrations palmelloid colony formation protects from MP-induced chlorophyll bleaching, membrane damage or oxidative stress.

**Table 1:**
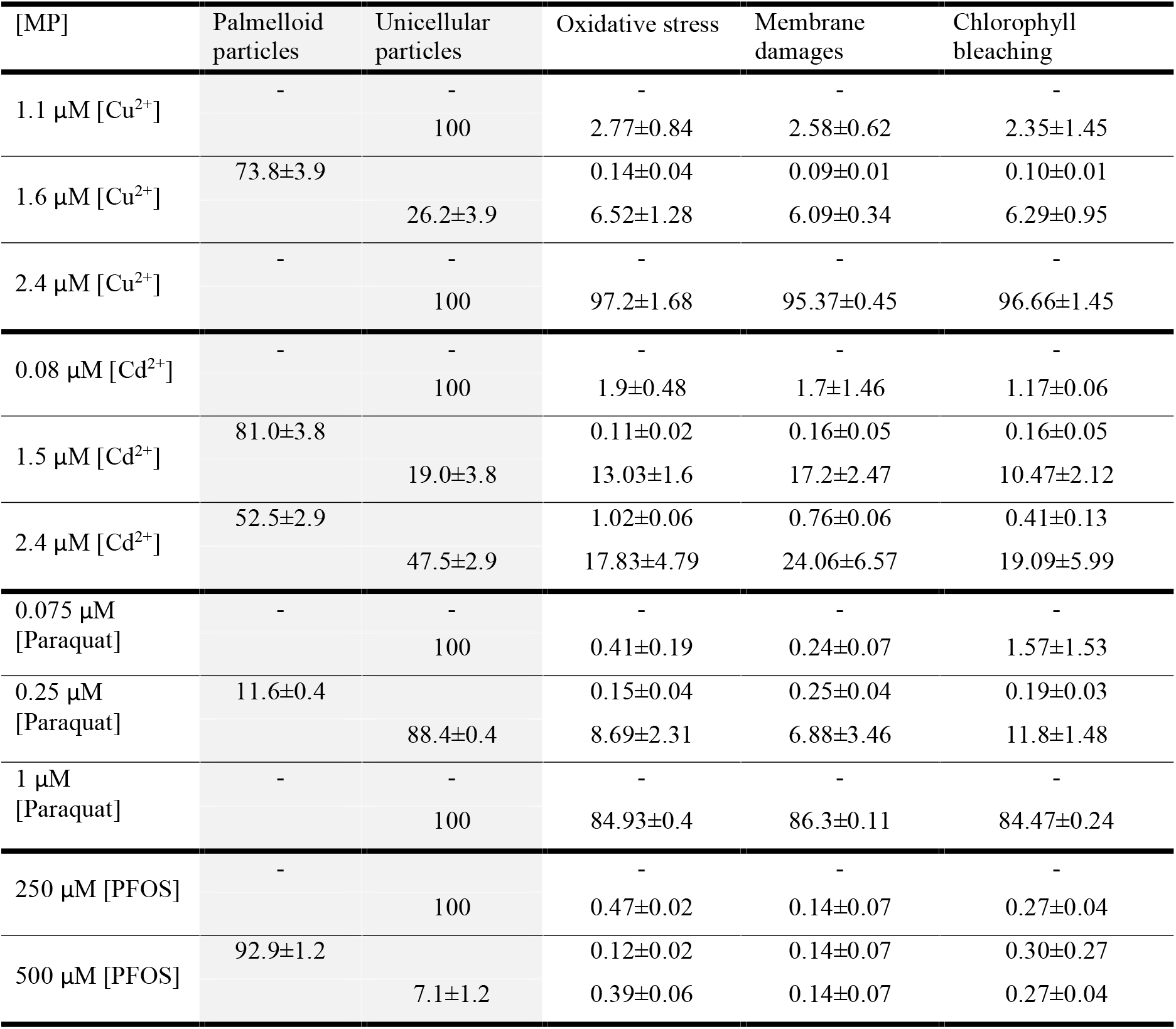
Percentage of unicellular or palmelloid particles affected by oxidative stress, membrane damages or chlorophyll bleaching after 48h exposure to Cu, Cd, PFOS and paraquat (mean values ± standard deviation).

### Palmelloid formation and Cadmium uptake

Cellular and colony morphology are important drivers of phytoplankton community assembly. Phytoplankton size and shape are known to play a determinant role in floating/sedimentation, grazing, light harvesting and nutrients availability (Naselli-Flores and Barone, 2011). Colony formation might have a cost for the organism since it is associated to higher rates of sedimentation that enhance the sinking out of the euphotic zone (Lurling and Van Donk, 2000). Palmelloid colonies obtained upon MPs exposure were found to lose their ability to perform phototaxis (Figure S3), likewise *Chlamydomonas* colonies obtained under predator pressure (Boyd et al., 2018) andwere also found to sediment quite fast (Figure S4). The larger colony size and associated lower surface to volumes ratio might be a disadvantage for nutrient availability but inversely it might become an advantage in the case of micropollutants exposure. We hypothesized that palmelloid colonies have lower MPs content than unicellular *C. reinhardtii* and that the cell wall that surrounds the colony might act as a barrier that prevents micropollutant uptake.

In order to verify this hypothesis we decided to focus on Cd uptake. Cd was selected among the MPs tested for two reasons: i) palmelloid formation with this metal was stable trough time; ii) Cd can be easily measured via ICP-MS. Cd concentration in *C. reinhardtii* particles at different time points during palmelloid formation was determined (Figure 6). Cd content of algal particles was found to increase considerably in the first 24 hours exposure and continued to increase even after palmelloids were formed. The size of palmelloid colonies formed upon Cd exposure significantly increase as a function of incubation time (Figure 4). Even if the amount of Cd per palmelloid particle increased, Cd content per particle bio-volume was found to be stable. These results suggest that even if Cd is taken up in the colony its concentration within the palmelloid particle volume remains constant due to the algal cell division and increase in the number of cells that form the palmelloid. In order to determine whether the amount of Cd taken up by palmelloid colonies is proportional the colony size, palmelloid colonies previously obtained upon exposure to 500 μM [PFOS] for 24, 48 and 72h were exposed to 1.5 μM [Cd^2+^] for 2 hours. Moles of Cd taken up by the colony increased with increasing palmelloid size, conversely the moles of Cd per particle biovolume decreased significantly.

**Figure 6:**
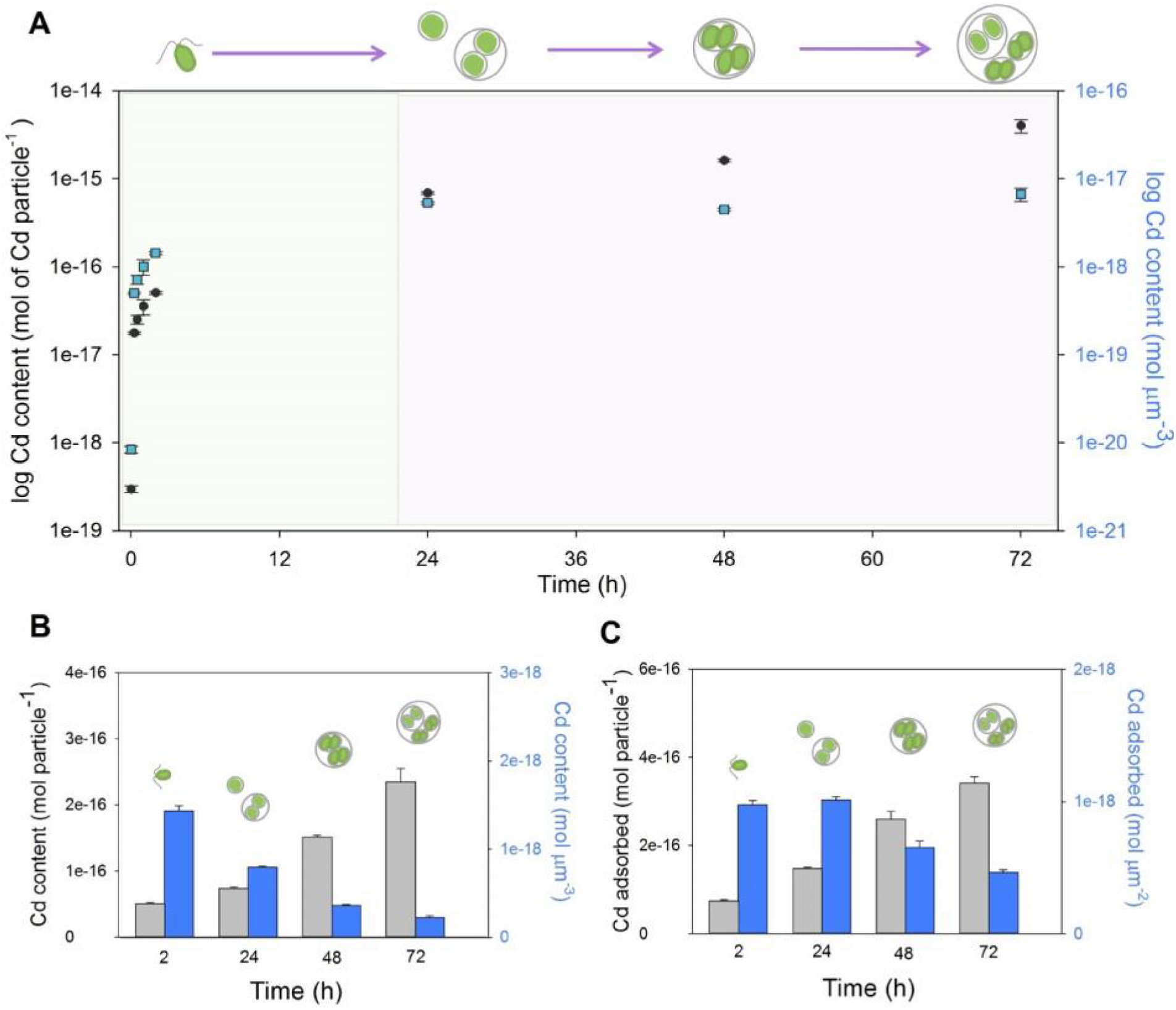
Cadmium content in unicellular *C. reinhardtii* and palmelloid colonies. A) Cd content as a function of exposure time during palmelloid formation upon exposure to 1.5 μM [Cd^2+^]. Black circles indicate moles of Cd per algal particle while blue squares indicate moles of Cd per particle volume. Results are the average of at least three independent replicates, error bars represent standard deviations. B) Cd content and C) Cd adsorbed after 2h exposure to 1.5 μM [Cd^2+^] of unicellular *C. reinhardtii* and palmelloid colonies of different size obtained upon exposure to PFOS for 24, 48 and 72h. Grey bars indicate moles per algal particle while blue bars indicate moles per particle volume (B) or particle surface (C).

The present results indicate that the cell wall surrounding palmelloid colonies is not a barrier that prevents metal uptake. However, the amount of Cd adsorbed per surface unit is lower in large palmelloid colonies than in small ones or in unicellular *C. reinhardtii*. Consequently lower Cd concentrations are measured in large palmelloid colonies with respect to small *C. reinhardtii* particles. *C. reinhardtii* cell walls are known to be positively involved in metal adsorption and uptake (Kola et al., 2004), higher intracellular Cd content was measured in a WT strain with respect to a cell wall-less strain (Kola and Wilkinson, 2005).

Overall data showed that palmelloid colonies adsorb lower Cd per surface units than unicellular *C. reinhardtii* and that the largest is the palmelloid colony the lower is its Cd concentrations, suggesting that the cells within large colonies experience lower Cd exposure than unicellular *C. reinhardtii*. What is more, the localization of Cd within the palmelloid particle remains to be elucidated. Nothing is known about the chemical composition of the cell wall that envelops palmelloid colonies and even less about the chemical composition of the volume that surrounds cells within the palmelloid colony. The EDTA non-extractable Cd fraction measured in the palmelloid might be not bioavailable for *C. reinhardtii* cells. In fact, it might be accumulated in the extracellular volume present within the palmelloid, adsorbed on the multiple cell wall layers or sequestrated inside different cellular compartments. Indeed, several detoxification mechanisms were described in *Chlamydomonas* to reduce toxic effects of elevated intracellular Cd quotas (Lavoie et al., 2009).

### Conclusions

The present study has demonstrated that the exposure to sub-lethal concentration of micropollutants with different modes of action resulted in the formation of palmelloid colonies. The palmelloid colony formation was a transitory response which was independent of the mode of MP action. Under these conditions, morphological changes are observed but not adverse toxic effects, indicating that such morphological plasticity is associated to stress acclimation responses and can be triggered to respond to micropollutant bursts. Investigation of processes associated to sudden exposure to stressful conditions are of particular relevance in order to understand how cells respond and acclimate to continuously changing environment. Morphological plasticity responses to MPs are expected to have indirect effects on phytoplankton dynamics and food webs. The investigation of species-level morphological plasticity are of primary importance for the prediction of community responses to MP stress.

## ASSOCIATED CONTENT

### Supporting Information

Table S1 reports the percentage of cells observed for each morphotype for all of the tested concentration; Figure S1 shows the gating strategy applied for the FCM analysis of MPs effects; Figure S2 presents the changes of dissolved total [Cu] in the exposure medium; Figure S3 shows phototaxis test in unicellular *C. reinhardtii* and palmelloid colonies; Figure S4 reports sinking behavior of palmelloid colonies.

The material is available free of charge via the Internet at

### Funding Sources

Giulia Cheloni acknowledges the Swiss National Science Foundation (SNF) for financial support (Marie Heim Vögtlin grants number PMPDP3_164428 and PMPDP_1837781).

## ACKNOWLEDGMENT

Giulia Cheloni acknowledge Isabel Worms for the support in ICP-MS data analysis, Séverine Le Faucheur for scientific discussion and Michel Goldschmidt-Clermont for scientific discussion and critical reading of the manuscript.

## ABBREVIATIONS

FCM: flow cytometry
MP: micropollutant
PFOS: Perfluorooctanesulfonic acid

