## Supplementary material for "Morphological plasticity in *Chlamydomonas reinhardtii* and acclimation to micropollutant stress": Table S1; Figure S1; Figure S2; Figure S3; Figure S4

Department F.-A. Forel for environmental and aquatic sciences, Earth and Environmental Sciences, Faculty of Sciences, University of Geneva, CH-1211 Geneva, Switzerland

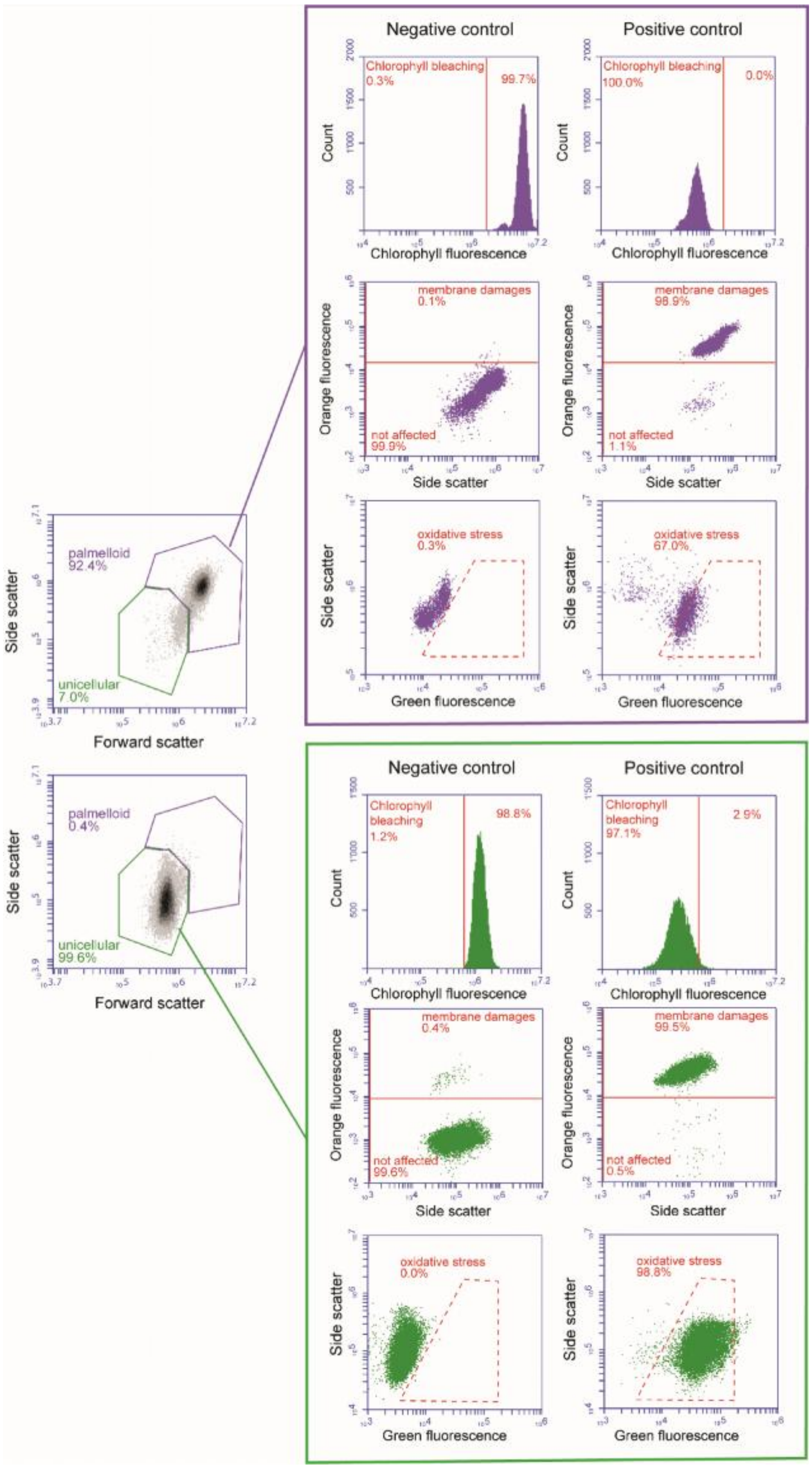


**Figure S1:** Gating strategy applied in the flow cytometric analysis adopted to evaluate toxic effects of MPs to algal particles. Unicellular *C. reinhardtii* were discriminated from palmelloid colonies by designing two gates that take advantage of the differences in side scattering and forward scattering properties of the algal particles. Specific gates were designed for each of the two populations in order to evaluate effects on chlorophyll bleaching, membrane damage and oxidative stress. The position of the gates was established taking into account the results obtained with positive control cultures. Positive controls of unicellular and palmelloid populations were obtained via exposure to UV light, 70°C temperature and cumene hydroperoxide for the endpoints of chlorophyll bleaching, membrane damage and oxidative stress respectively.

**Table S1**: Percentages of cells observed for each morphotype under different treatments.

| | [MPs] (µM) | Cells in palmelloids | | | Unicellular | | | Unicellular damaged | | |
|---|---|---|---|---|---|---|---|---|---|---|
| | | 24h | 48h | 72h | 24h | 48h | 72h | 24h | 48h | 72h |
| Cu | 0 | 0 | 0 | 0 | 100 | 100 | 100 | 0 | 0 | 0 |
| | 0.1 | 0 | 0 | 0 | 100 | 100 | 100 | 0 | 0 | 0 |
| | 0.4 | 0 | 0 | 0 | 100 | 100 | 100 | 0 | 0 | 0 |
| | 0.7 | 0 | 0 | 0 | 100 | 100 | 100 | 0 | 0 | 0 |
| | 1.1 | 0 | 0 | 0 | 100 | 100 | 100 | 0 | 0 | 0 |
| | 1.3 | 20.6±2.4 | 2.4±3.1 | 78.2±3.2 | 79.4±2.4 | 21.8±3.1 | 75.4±3.2 | 0 | 0 | 0 |
| | 1.6 | 26.8±2.9 | 2.9±3.9 | 87.8±1.1 | 73.2±2.9 | 12.2±3.9 | 60.6±1.1 | 0 | 0 | 0 |
| | 1.9 | 0 | 0 | 0 | 100 | 89.2±2.1 | 68.1±3.8 | 0 | 10.8±2.1 | 31.9±3.8 |
| | 2.2 | 0 | 0 | 0 | 100 | 78.3±3.5 | 81.8±10 | 0 | 21.7±3.5 | 18.2±10 |
| | 2.5 | 0 | 0 | 0 | 100 | 79.9±5.0 | 64.9±5.6 | 0 | 20.1±5.0 | 35.1±5.6 |
| Cd | 0 | 0 | 0 | 0 | 100 | 100 | 100 | 0 | 0 | 0 |
| | 0.01 | 0 | 0 | 0 | 100 | 100 | 100 | 0 | 0 | 0 |
| | 0.08 | 0 | 0 | 0 | 100 | 100 | 100 | 0 | 0 | 0 |
| | 0.4 | 13.6±6.0 | 46.6±5.4 | 41.7±5.9 | 86.4±6.0 | 51.7±7.0 | 76.9±6.1 | 0 | 1.7±1.5 | 1.2±0.3 |
| | 0.7 | 38.0±7.1 | 53.0±3.9 | 58.6±7.2 | 72.6±2.1 | 46.0±3.6 | 57.1±1.2 | 0 | 1.0±0.4 | 3.4±0.9 |
| | 1.1 | 17.9±4.5 | 81.5±3.9 | 51.0±1.1 | 82.1±4.5 | 18.5±3.9 | 47.3±1.5 | 0 | 0 | 1.8±0.4 |
| | 1.5 | 24.0±1.5 | 78.0±3.8 | 86.2±1.1 | 76.0±1.5 | 19.0±3.4 | 11.4±0.9 | 1.1±0.3 | 3.0±0.5 | 2.4±0.2 |
| | 1.9 | 44.3±1.4 | 80.4±1.4 | 89.4±0.8 | 55.7±1.4 | 16.5±1.3 | 8.2±0.5 | 8.7±1.2 | 3.0±0.1 | 2.4±0.2 |
| | 2.4 | 39.5±8.7 | 52.5±2.9 | 58.2±3.1 | 51.5±5.0 | 37.1±1.4 | 20.8±0.6 | 9.0±3.7 | 10.4±4.3 | 21.0±3.6 |
| PFOS | 0 | 0 | 0 | 0 | 100 | 0 | 100 | 0 | 0 | 0 |
| | 50 | 0 | 0 | 0 | 100 | 0 | 100 | 0 | 0 | 0 |
| | 100 | 0 | 0 | 0 | 100 | 0 | 100 | 0 | 0 | 0 |
| | 200 | 0 | 0 | 0 | 100 | 0 | 100 | 0 | 0 | 0 |
| | 250 | 0 | 0 | 0 | 100 | 0 | 100 | 0 | 0 | 0 |
| | 300 | 15.3±3.1 | 17.1±1.9 | 0.5±0.7 | 84.7±3.1 | 82.9±1.9 | 99.5±0.7 | 0 | 0 | 0 |
| | 350 | 22.2±2.5 | 13.4±2.0 | 2.4±0.7 | 77.8±2.5 | 86.6±2.0 | 97.6±0.7 | 0 | 0 | 0 |
| | 500 | 56.6±3.7 | 97.9±1.2 | 98.9±0.1 | 43.4±3.7 | 2.1±1.2 | 1.1±0.1 | 0 | 0 | 0 |
| | 750 | 93.8±1.1 | 99.2±0.1 | 99.8±0.3 | 6.2±1.1 | 0.8±0.1 | 0.2±0.3 | 0 | 0 | 0 |
| Paraquat | 0 | 0 | 0 | 0 | 100 | 0 | 100 | 0 | 0 | 0 |
| | 0.025 | 0 | 0 | 0 | 100 | 0 | 100 | 0 | 0 | 0 |
| | 0.05 | 0 | 0 | 0 | 100 | 0 | 100 | 0 | 0 | 0 |
| | 0.075 | 0 | 0 | 0 | 100 | 0 | 100 | 0 | 0 | 0 |
| | 0.1 | 7.4±1.1 | 9.1±3.2 | 7.3±2.8 | 92.6±1.1 | 90.9±3.2 | 92.7±2.8 | 0 | 0 | 0 |
| | 0.25 | 6.7±0.2 | 18.6±0.4 | 37.2±5.3 | 93.3±0.2 | 81.4±0.4 | 62.8±5.3 | 0 | 0 | 0 |
| | 0.5 | 3.4±0.7 | 12.5±3.0 | 24.4±0.6 | 96.6±0.7 | 87.5±3.0 | 62.9±0.5 | 0 | 0 | 12.7±1.1 |
| | 0.75 | 9.4±4.1 | 12.2±0.6 | 6.9±2.7 | 90.6±4.1 | 87.8±0.6 | 56.4±0.7 | 0 | 0 | 36.8±3.4 |
| | 1 | 0 | 0 | 0 | 100 | 96.5±0.9 | 100 | 0 | 3.5±0.9 | 87.6±5.8 |
| | 2.5 | 0 | 0 | 0 | 100 | 91.8±2.6 | 100 | 0 | 8.2±2.6 | 82.3±6 |

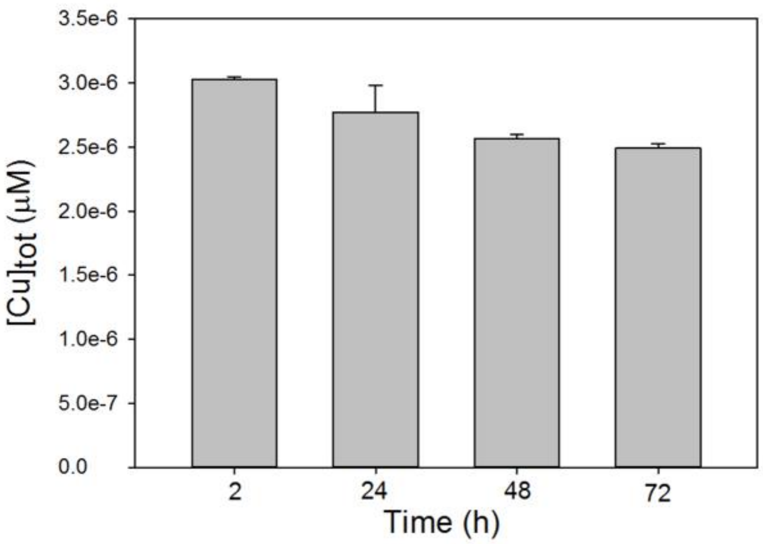


**Figure S2**: Total dissolved [Cu] measured in cultures exposed to 1.6 µM [$Cu^{2+}$]

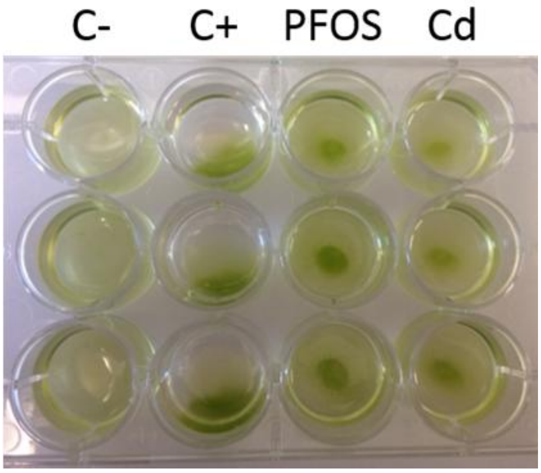


**Figure S3**: Phototaxis test performed on unicellular *C. reinhardtii* (C+). Temperature inactivated unicellular *C. reinhardtii* (C-) and palmelloid colonies obtained after 48h exposure to 500 uM PFOS or 2.6 uM Cd. Plates were incubated for 30 minutes under a unidirectional light source (cool white lamp, 30 mmol phot $m^{-2}$ $s^{-1}$.

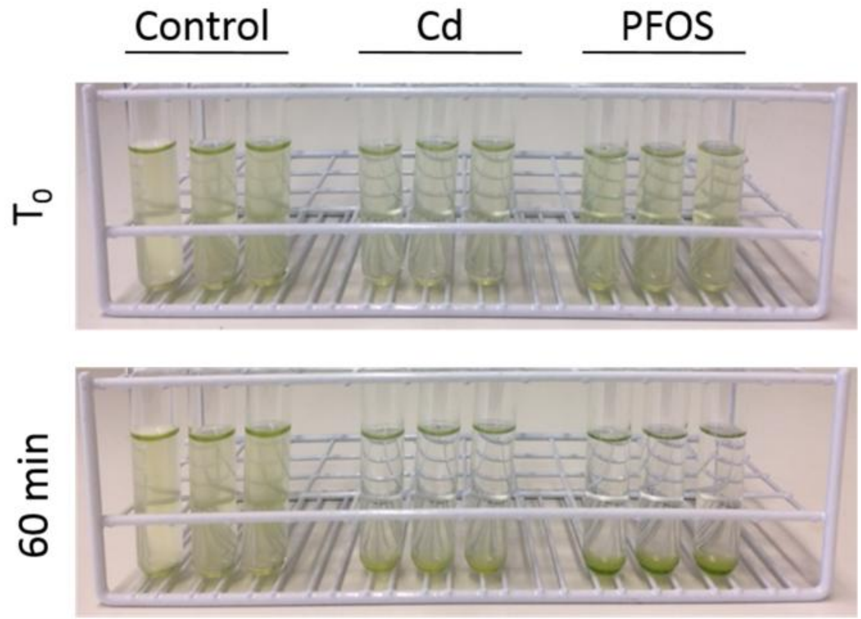


**Figure S4**: Sinking behaviour of unicellular *C. reinhardtii* cells and palmelloid colonies.
